# Belief embodiment through eye movements facilitates memory-guided navigation

**DOI:** 10.1101/2023.08.21.554107

**Authors:** Akis Stavropoulos, Kaushik J. Lakshminarasimhan, Dora E. Angelaki

**Author notes:** Correspondence: Dr. Akis Stavropoulos Center for Neural Science, Mayer 901 New York University, NY 10003. Equal author contribution (Akis Stavropoulos, Kaushik J. Lakshminarasimhan).

## Abstract

Neural network models optimized for task performance often excel at predicting neural activity but do not explain other properties such as the distributed representation across functionally distinct areas. Distributed representations may arise from animals’ strategies for resource utilization, however, fixation-based paradigms deprive animals of a vital resource: eye movements. During a naturalistic task in which humans use a joystick to steer and catch flashing fireflies in a virtual environment lacking position cues, subjects physically track the latent task variable with their gaze. We show this strategy to be true also during an inertial version of the task in the absence of optic flow and demonstrate that these task-relevant eye movements reflect an embodiment of the subjects’ dynamically evolving internal beliefs about the goal. A neural network model with tuned recurrent connectivity between oculomotor and evidence-integrating frontoparietal circuits accounted for this behavioral strategy. Critically, this model better explained neural data from monkeys’ posterior parietal cortex compared to task-optimized models unconstrained by such an oculomotor-based cognitive strategy. These results highlight the importance of unconstrained movement in working memory computations and establish a functional significance of oculomotor signals for evidence-integration and navigation computations via embodied cognition.

## Introduction

The brain evolved complex recurrent networks to interpret and act upon a dynamic and uncertain world, but its computational powers and mechanisms generating natural behavior remain cryptic (Cooper & Peebles, 2015; Krakauer et al., 2017; Miller et al., 2022). Most of our insights into neural computation are based on binary tasks with highly constrained actions that are artificially segregated from perception (Gold & Shadlen, 2007; Freedman & Assad, 2011; Murray et al., 2014; O’Connell et al., 2018). Tightly controlling laboratory behavior by preventing natural, continuous movements has simplified interpretability but also hindered our ability to gain insights from natural behavioral strategies. Artificially keeping the eyes fixed, for example, has been standard in monkey studies of working memory and decision-making (Cohen & Maunsell, 2011; Kiani & Shadlen, 2009; Meirhaeghe et al., 2021; O’Shea et al., 2004; Ruff & Cohen, 2014). In contrast, natural behavior involves continuous eye movements (Hayhoe & Ballard, 2005; Hayhoe et al., 2012; Mao et al., 2021). Thus, there is concern that traditional experimental paradigms, such as those requiring fixation or tightly constrained actions, deprive subjects of natural affordances (Gibson et al., 1978), and thus may hide fundamental neural mechanisms that are only expressed in a dynamic, closed-loop context. Can more naturalistic laboratory behaviors with free eye movements shed new light onto neural mechanisms of ethological behaviors?

A major gap in understanding is epitomized by an emerging tool to probe neural mechanisms: neural network models optimized to perform neuroscience tasks. The representations learned by the networks often resemble the response properties of brain areas that drive behavior in those tasks (Yamins & DiCarlo, 2016; G. R. Yang et al., 2019). However, such models are typically grounded in generic neural architectures – feedforward or recurrent depending on the task – and cannot explain why neural computations are distributed across functionally distinct circuits. A jarring example of distributed brain computation is the prevalence of motor signals in sensory and association areas (Hadjidimitrakis et al., 2019; Musall, Kaufman, et al., 2019), and sensory signals in the motor and frontal areas (Ebbesen et al., 2018; Zaksas & Pasternak, 2006). Existing models of distributed neural representations appeal to the brain’s recurrent architecture to capture multi-area data but fall short of providing a normative account of such representations (Kleinman et al., 2021; G. R. Yang & Molano-Mazón, 2021). There is a growing realization that building task-optimized neural network models with brain-inspired modular architectures provide limited insights beyond what is already determined by the task goal (Michaels et al., 2020; Pinto et al., 2019). To gain new insights from adopting brain-like architectures, we need to additionally incorporate the specific strategy used by animals to solve the task (Musall, Urai, et al., 2019). Traditional neuroscience tasks like binary decision-making are too simple to admit interesting cognitive strategies, especially when participants are mechanically restrained in fixation-based paradigms.

To unravel the neural mechanisms of natural behavior and to interrogate alternatives to the traditional approach, we have developed a naturalistic navigation task featuring action/perception loops in virtual reality (VR) with unconstrained eye movements. This continuous foraging task requires participants to steer towards remembered target locations by using sensory evidence, working memory, and continuous actions constituting a naturalistic visual perception-action loop (Lakshminarasimhan et al., 2018, 2020). Participants observe a briefly flashed target in the distance (like the blinking of a firefly) and steer to the remembered target location using optic flow feedback from a virtual environment comprising an unstructured ground plane with no landmarks. Importantly, in contrast to traditional tasks such as evidence accumulation or delayed discrimination in which the latent world states and/or contents of working memory remain unchanged throughout the trial, the latent state (i.e., egocentric target location) dynamically varies over the course of each trial, under the participant’s control, and must be mentally tracked in order to know precisely when to stop steering.

In principle, this task can be performed without physically tracking the believed goal location with one’s eyes. Yet, Lakshminarasimhan et al. (2020) found that both humans and monkeys tend to follow the location of the invisible target with their gaze until they reach it and noticed a significant decline in steering performance when eye movements were suppressed. Given the visually guided nature of the steering task, such eye movements may reflect a strategy to gather information about self-motion: since subjects must integrate optic flow to dynamically update their beliefs about the relative goal location, directing gaze to specific regions of the environment such as the focus of expansion, might help acquire more information about their movement velocity (*active sensing* hypothesis). Alternatively, these task-relevant eye movements may reflect an embodiment of subjects’ dynamically evolving internal beliefs about the goal: by allowing dynamic beliefs about the relative target location to continuously modulate eye movements in this task, the brain could piggyback on the oculomotor circuit and reduce the computational burden on working memory (*cognitive embodiment* hypothesis). The latter hypothesis predicts that these eye movements should also govern other types of navigation, e.g., inertially guided steering, where the joystick controls inertial accelerations in the absence of visual cues.

Here we first provide strong support for the embodiment hypothesis by analyzing eye movements under both inertially and visually guided versions of the task. We found that under both sensory conditions eye movements reflect the evolving belief dynamics about the relative target location. We then used this behavioral strategy as an additional constraint for training a distributed neural network model and found that it recapitulated the behavioral and neural data more accurately with fewer tunable parameters than purely task-optimized models. These results lend support to the notion that ethologically valid paradigms can help constrain modeling to provide new insights into the neural mechanisms and the emergence of distributed neural representations.

## Results

Participants sitting on a motion platform performed a virtual reality (VR) navigation task using a joystick to steer freely and catch targets that pop up transiently (like fireflies) on the ground plane, one at a time (**Fig. 1A**). Participants’ steering was coupled either to a visual environment that provided optic flow but devoid of landmarks (‘Visual’ condition) or to the platform’s motion in complete darkness (‘Inertial’ condition; for details see Stavropoulos et al., 2022). At the beginning of each trial, a target (‘firefly’) appears for 1s at a random location on the ground plane within the field of view (**Fig. 1B**). When it disappears, an analog joystick controlling linear and angular motion is activated allowing participants to navigate toward the remembered target location. Participants could steer freely on the ground plane and integrate momentary sensory evidence about their movements based on either visual (optic flow) or inertial (vestibular with somatosensory) sensory cues. This task has a crucial time-varying latent variable: position of the target relative to oneself, which must be computed by integrating noisy angular/linear sensory cues (visual or inertial), which in turn are controlled dynamically by the participant’s joystick actions. A time constant governed the control dynamics: in trials with a *small* time constant, joystick position mainly controlled velocity (Velocity Control; VC); when the time constant was *large*, joystick position mainly controlled acceleration (Acceleration Control; AC), mimicking inertia under viscous damping (**Fig. 1C**). Across trials, visual and inertial sensory conditions were randomly interleaved while manipulation of the time constant followed a bounded random walk (Stavropoulos et al., 2022).

**Figure 1:**
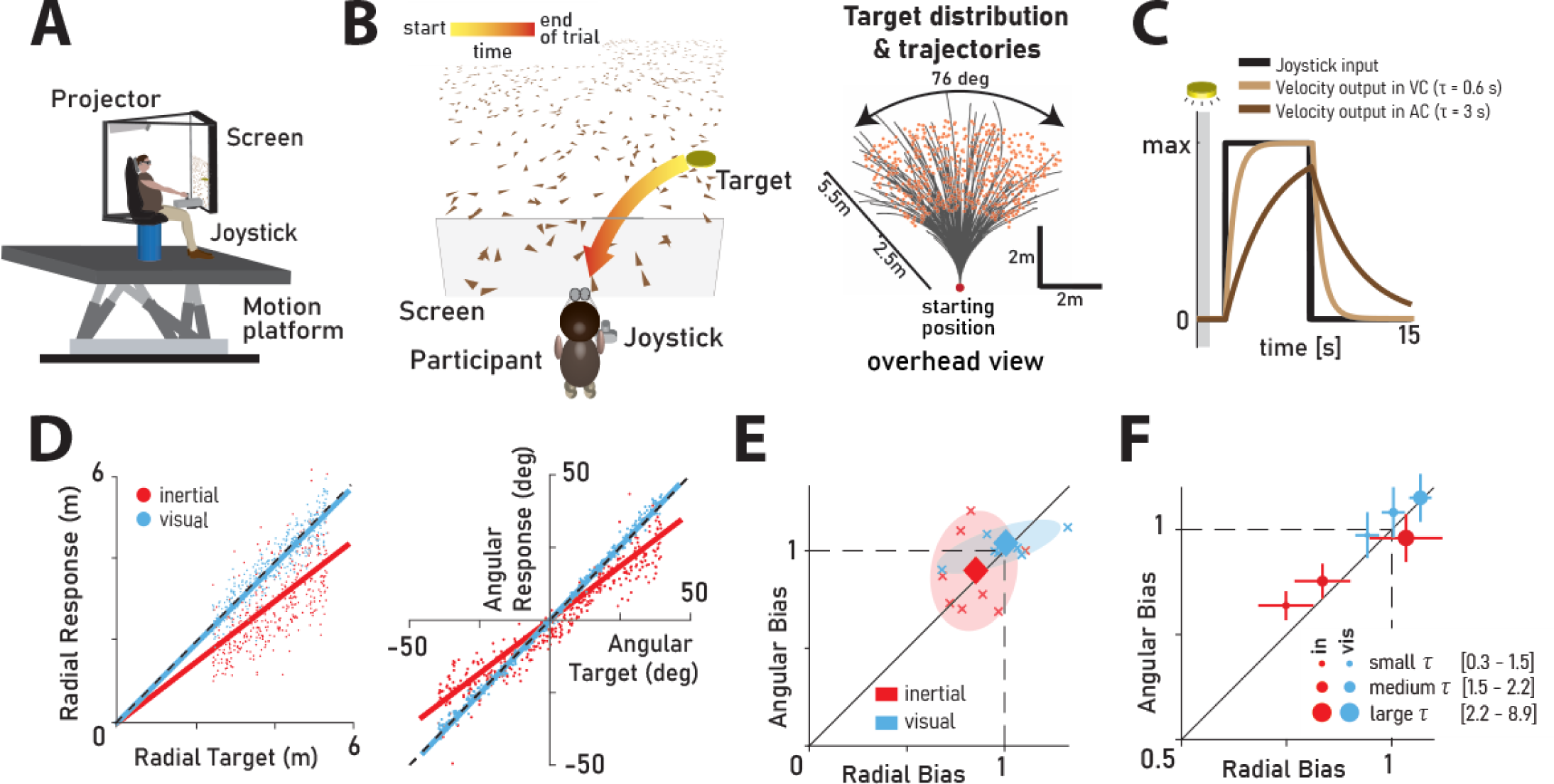
Experimental design and task performance. **(A)** Experimental set up. **(B) Left:** Illustration of virtual environment. Participants steer towards a briefly cued target (*yellow disc*) using optic flow cues available on the ground plane (visual condition only; platform motion is the only available cue in the inertial condition). During steering, the target becomes less eccentric over time (towards participant’s midline), while it lowers in the participant’s field of view (*color-coded arrow*). **Right:** Overhead view of the spatial distribution of target positions across trials and the corresponding trial trajectories. *Red dot* shows the starting position of the participant. **(C)** Simulated maximum pulse joystick input and the corresponding velocity output under Velocity Control (VC; *beige*) and Acceleration Control (AC; *brown*). The input is lowpass filtered to mimic the existence of inertia. The time constant of the filter varies across trials (time constant *τ*), along with maximum velocity to ensure comparable travel times across trials. Gray zone: brief cueing period of the target at the beginning of the trial. **(D)** Target vs Response. **Left:** Comparison of the radial distance of a typical subject (stopping location) against radial distance of the target across all trials. **Right:** Angular eccentricity of the response vs target angle. Black dashed lines have unity slope (unbiased performance). Solid lines: linear regression. Data colored according to the sensory condition *(red:* inertial, *cyan:* visual). Radial and angular response biases were defined as the slope of the corresponding regressions. **(E)** Scatter plot of radial and angular biases in each sensory condition plotted for each individual participant. *Ellipses* show 68% confidence intervals of the distribution of data points for the corresponding sensory condition. *Diamonds* (centers of the ellipses) represent the mean radial and angular response biases across participants. Dashed lines indicate unbiased radial or angular position responses. Solid diagonal line has unit slope. **(F)** Participant average of radial and angular response biases in each condition, with trials grouped into tertiles of increasing time constant *τ*. Error bars denote ±1 SEM.

Subjects did not receive any performance-related feedback. By varying motion dynamics across trials and by eliminating performance feedback, we aimed to induce variable behavioral performance which ensures greater statistical power in the analyses needed to decouple subjective beliefs from the true latent states. Indeed, subjects showed strong biases, especially in the inertial condition (Fig. 1D, E). Biases, defined as the regression slope between target and stopping positions (a value of 1 indicates no bias), are strongly correlated with the control dynamics (Fig. 1F), a pattern that has been described in detail previously (Stavropoulos et al., 2022). Here we compare the eye movements generated in the inertial, compared to the visual version of the task, to distinguish between the *active sensing* and *cognitive embodiment* hypotheses.

### Eye movements track beliefs about the latent goal location

Participants received no instruction about their gaze behavior, yet eye movements tracked the memorized location of the goal, and this was true not only in the visual condition (as previously shown by Lakshminarasimhan et al., 2020), but also during inertially-guided steering. As the target is on the ground plane below subjects’ eye level (Fig. 1A**-B**), the relative position of the (invisible) target approaches the midline and moves downward in the visual field as the participant steers towards it. Eye movements mirrored this same pattern: horizontal eye position converged toward zero (midline) and vertical eye position descended over time, even under the Inertial condition, where no visual cues were provided after the target disappeared (Fig. 2A). As shown in a typical example, there was an initial saccade towards the target immediately after target onset (Fig. 2B – gray region, 0-1s), followed by a mostly smooth tracking until the end of the trial.

**Figure 2:**
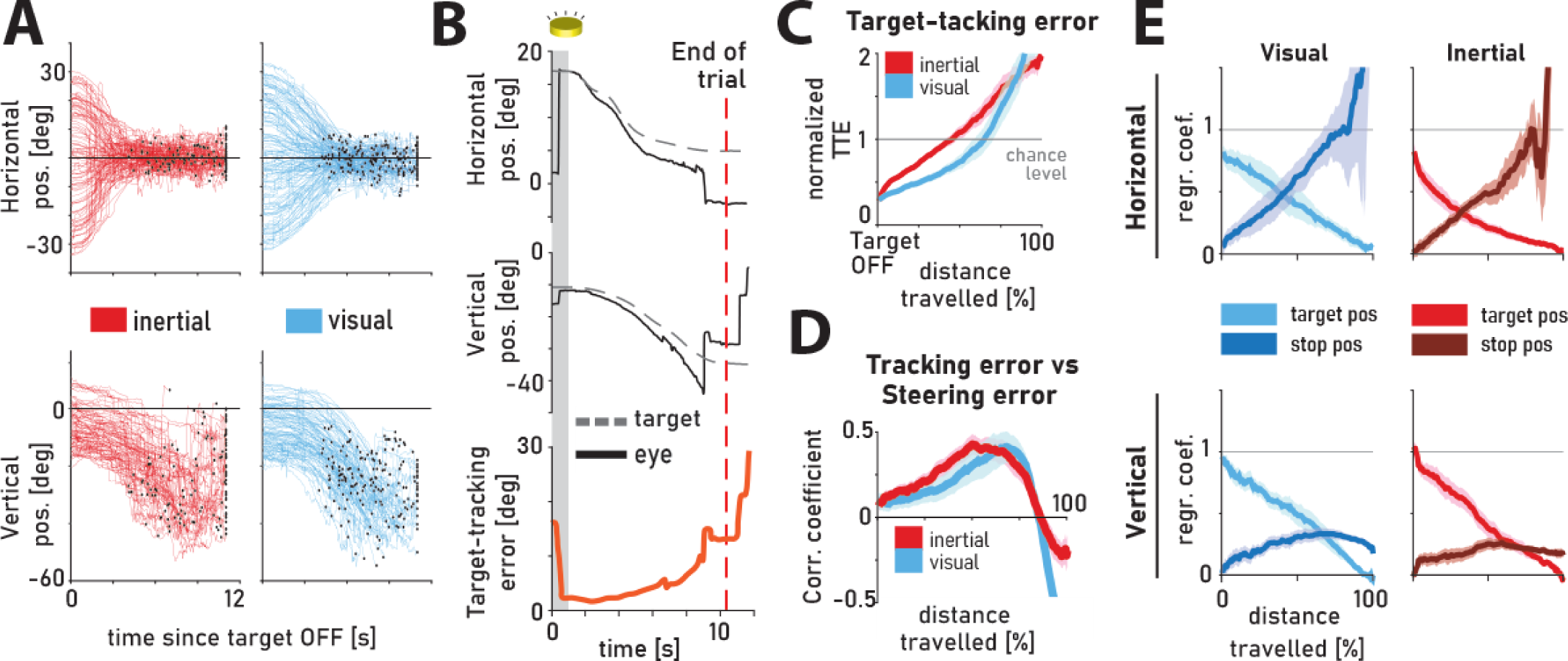
Target-tracking with eye movements. **(A)** Time-course of horizontal and vertical eye position during a random subset of trials from one participant; time at 0 denotes disappearance of target (target offset); Black dots illustrate end of trial (clipped at 11s). **(B)** Time-course of horizontal (*top*) and vertical (*middle*) eye position (*black solid lines*), the respective target position (*gray dashed lines*), as well as target-tracking error (TTE, *orange*), during a representative trial. Gray region denotes the period when a target was visible. Red dashed line corresponds to the end of the trial. **(C)** Normalized TTE over trial progression (percentage of total distance travelled). TTE was normalized by the chance-level TTE obtained by shuffling (*gray line*). The area above the gray line corresponds to TTE worse than chance. *Shaded regions:* ±1 SEM across subjects. **(D)** Correlation coefficient of TTE and steering error (SE). *Shaded regions:* ± 1 SEM across participants. **(E)** Multilinear regression of eye position against initial target and stop positions over trial progression, for both horizontal and vertical components. *Light* and *dark shades* represent the target and stop position coefficients, respectively. Notice how the modulation of eye position by the target and stop positions gradually reverses as the trial progresses. *Shaded regions:* ± 1 SEM across participants.

We computed the Euclidean distance between *eye* and *target* position as the “target-tracking error (TTE)” (**Fig 2B**, bottom, see **Methods**). TTE at target offset was low across subjects (mean TTE at trial onset ± standard deviation: 5.60 ± 0.38°) and increased as the trial progressed (Fig. 2C). Despite the extremely long trial durations (across subjects trial duration mean ± standard deviation – inertial: 14.1 ± 5.1 s, visual: 13.3 ± 4 s), TTE remained significantly below chance level obtained by shuffling (**Methods**; Fig. 2C, *gray line*) for 68.8±4.9 % (visual) and 51.9±4.4 % (inertial) of the trajectory (mean ± SEM of percentage distance travelled until TTE crosses chance level; for data from individual subjects see **Suppl. Fig. S1A**). These results are consistent and build upon findings from a purely velocity-controlled visual steering task of much shorter trial durations (∼2 s) with performance feedback (Lakshminarasimhan et al. 2020). Notably, these results hold true also for inertial navigation in the absence of optic flow, suggesting that the pattern of eye movements reflects a strategy of embodiment and is not linked solely to active sensing of optic flow patterns, if at all. While TTE was larger for the Inertial condition compared to the Visual condition (Fig. 2C, red vs. blue), this is due to increased behavioral variability in the former condition (**Suppl. Fig. S1B**) rather than an inability to track the memorized goal.

A trivial explanation for the increase in TTE over time is that eye movements become progressively more random with time. Alternatively, the increase in TTE could arise if eye movements track the participant’s *belief* about the goal location rather than the true goal location. In this case, TTE should correlate with steering error (SE, *distance of stopping from actual target position*). This is because steering decisions are based on beliefs, and consequently any error in belief must manifest as an error in the eventual stopping position. Indeed, the correlation between TTE and SE increased as the trial progressed, reaching a peak at about 70% (visual) and 50% (inertial) into the trial, and decreased sharply thereafter (Fig. 2D). Peak correlations were statistically significant (p<0.05) individually in almost all participants (inertial: 8/8 participants, visual: 7/8). This result supports the hypothesis that the eyes track the *believed* location of the target in the virtual environment. In fact, when regressed against both the initial target and stop positions, eye movements were driven mostly by target position at trial onset and mostly by stopping position at the end of the trial (Fig. 2E), revealing how believed target location drifts gradually from the target to the stop location over the course of a trial.

### Small saccades aid belief tracking

During steering, saccade frequency (**Suppl.** Fig. 2A**, top**) and amplitude (**Suppl.** Fig. 2A**, bottom**) were both suppressed. Nevertheless, the infrequent small saccades contributed to goal tracking, as there was a drop in the correlation between steering and tracking error when saccades were removed from the data (Fig. 3A). This was particularly notable in the inertial condition, where the horizontal slow eye movements are strongly affected by the yaw vestibulo-ocular reflex (VOR; **Suppl.** Fig. 2B). Indeed, there were significant correlations between the cumulative horizontal saccade amplitude and the angular steering errors, revealing that saccades made during the VOR had a major contribution in ‘undoing’ the effects of the VOR such that the eye position could still reflect the internal belief of goal location (Fig. 3B). We previously have shown that forcing participants to fixate (in the visual condition) substantially affects task performance (Lakshminarasimhan et al. 2020). Combined with the persistent target-tracking in darkness while under the influence of VOR, these findings strongly suggest that this embodiment has a computational role.

**Figure 3:**
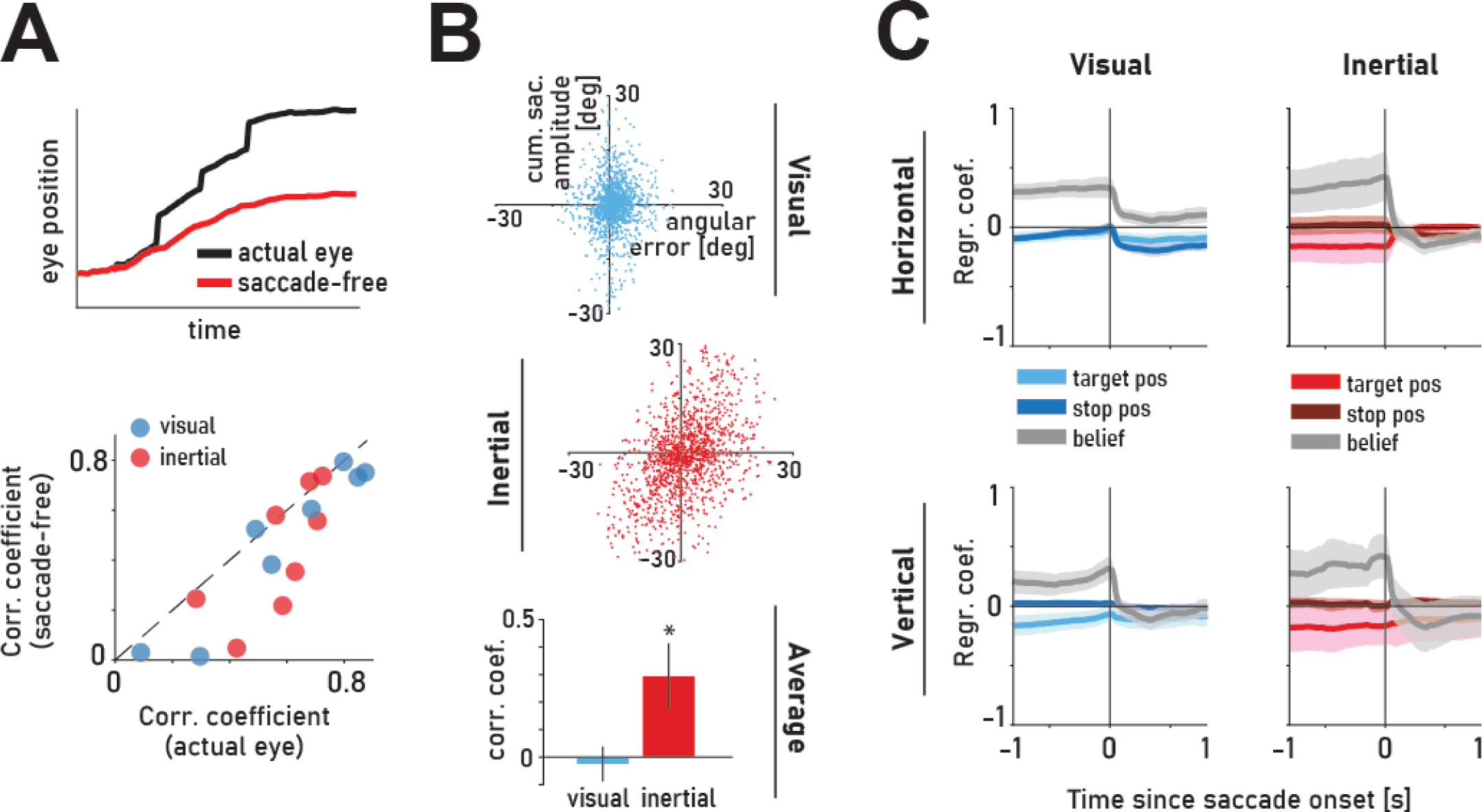
Saccades contribute to evolving belief about goal location. **(A) Top:** actual eye position (*black line*) and the corresponding saccade-free eye position (*red line*) of an example trial. **Bottom:** Comparison of correlation coefficients of target-tracking error (TTE) and steering error (SE) for the actual eye position and saccade-free eye position (horizontal component). Peak correlations within 50% of distance travelled were selected for each participant. **(B)** Correlation between cumulative horizontal saccade amplitude and angular steering error in the visual (**top, blue**) and inertial (**middle, red**) conditions. Saccades (within 50% of distance traveled) were pooled across participants for better visualization. **Bottom**: average correlation coefficients across participants. *Asterisks* denote the level of statistical significance of the correlation difference within each condition (*: p<0.05). Error bars: ±1 SEM. **(C)** Time-course (kernel) of coefficients obtained by linearly regressing the amplitude of the horizontal/vertical component of saccades evoked within 50% of distance travelled against the corresponding target-tracking error (*light blue/red*), stop position-tracking error (*dark blue/red*), and believed target-tracking position (*gray*). *Shaded regions* denote ± 1 SEM obtained by bootstrapping.

To more directly explore what drives saccadic eye movements during steering, we ran a regression analysis that shows how saccade amplitude is modulated by errors in tracking the actual target position, the stop position, or the participant’s reconstructed dynamic *belief* about goal location – the latter computed as the weighted average of the actual target and stop positions over time obtained for each participant (from Fig. 2E). Saccades are indeed modulated by beliefs, as illustrated by the fact that, before saccade-onset, the kernel (time-course of the coefficients obtained by linear regression) is larger for the tracking-error corresponding to the believed goal rather than the actual target or stop position (Fig. 3C; pre-saccadic peak of regression kernel mean ± SEM, horizontal component – inertial: [target: -0.05±0.02, stop: 0.03±0.07, belief: 0.43±0.21], visual: [target: -0.02±0.06, stop: - 0.002±0.05, belief: 0.34±0.09]; vertical component – inertial: [target: -0.16±0.16, stop: 0.05±0.03, belief: 0.42±0.19], visual: [target: -0.10±0.06, stop: 0.03±0.02, belief: 0.32±0.10). These results support the hypothesis that the small saccades generated during steering stabilize the gaze towards the believed goal location.

In summary, we conclude that participants integrate movement velocity to track their position relative to the goal using an oculomotor-based cognitive strategy: the evolving belief about goal location relative to their current position is embodied in eye position - and this cognitive embodiment has a computational role. We now turn to modeling to understand how such a strategy of embodiment can inform the underlying neural mechanisms. Specifically, we train different neural models optimized to do this task both with and without this cognitive strategy and evaluate how well each model predicts behavioral and neural data recorded in monkeys.

### A frontoparietal network model constrained by behavioral strategy

We previously demonstrated that both posterior parietal and dorsolateral prefrontal cortices represent latent beliefs in this task (Noel et al., 2022; Lakshminarasimhan et al., 2023). For simplicity, here we consider a combined frontoparietal recurrent neural network (FPN) as a stand-in for computations across both cortical areas. To investigate the mechanistic contribution of eye movements to this network, we simulated four models that are architecturally identical but differ in which connections are tuned (Fig. 4A – green and crimson). They feature a frontoparietal (FPN) module (see Discussion) comprising 100 recurrently connected nonlinear (‘sigmoidal’) units that receive two-dimensional sensory inputs (linear and angular velocity) and a two-dimensional pulse whose amplitude encodes the target position (x-y coordinates) at the beginning of each trial. The FPN module sends projections to motor units that drive two-dimensional joystick actions (linear and angular acceleration) and has bidirectional connections with oculomotor (OC) units that drive two-dimensional eye movements (horizontal and vertical). Two of the models are optimized solely for task performance i.e., to minimize the discrepancy between the stopping position and the target position, by tuning either just the readout weights onto the motor units (**Model 1**) or both the readout and recurrent weights within FPN (**Model 2**). The remaining models are optimized for task performance by tuning the readout weights, while also being constrained by the strategy used by humans and monkeys. Specifically, we tune the weights from FPN to OC to minimize an auxiliary loss such that OC could dynamically decode position relative to the target from FPN activity (**Models 3 & 4**). We additionally tune the feedback projection from OC to FPN to optimize for task performance (**Model 4**). For simplicity, we ignore recurrence within modules other than FPN. We trained the network weights (green weights) to reach the target in a time-bound manner via backpropagation-through-time. OC neurons in Models 3 & 4 are constrained to encode the relative target location via linear regression (crimson weights) (Fig. 4B). In all models, observation and process noise were added to the sensory and motor units respectively to prevent the models from developing a purely feedforward control strategy (**Methods**).

**Figure 4:**
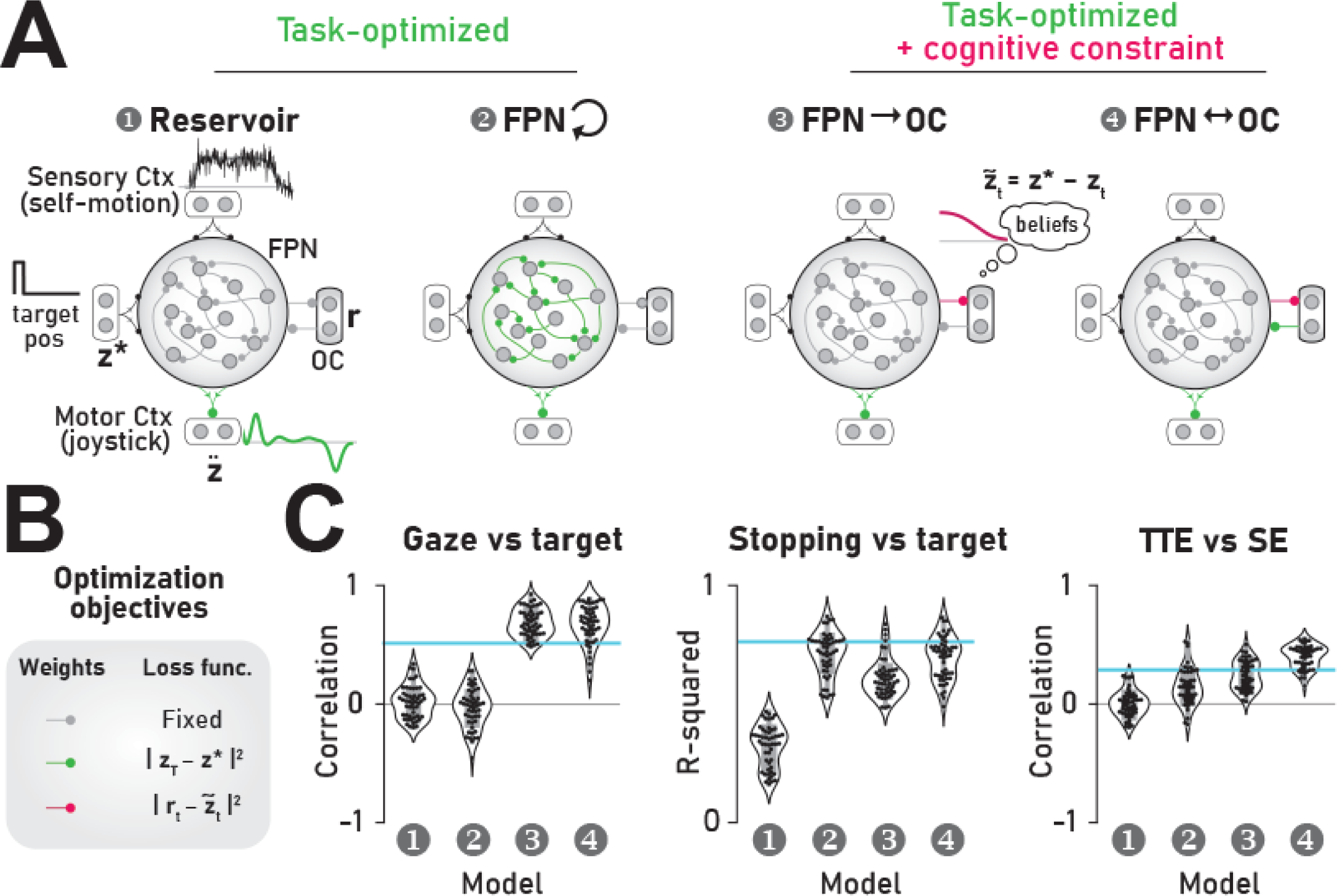
Model comparison. **(A)** Model Schematic. Models comprise a posterior parietal cortex (FPN, n=100 units) module which continuously receives 2D self-motion velocity and transiently receives 2D target position as inputs. PPC is recurrently connected with oculomotor (OC, n=2) module. Linear readout of FPN activity drives motor output which encodes 2D acceleration (linear/angular). For simplicity, the traces show only one dimension. **(B)** In all models, connectivity weights shown in gray are fixed, while weights in green are optimized for task performance by minimizing the squared error between stopping position (𝐳_T_) and target position (𝐳^∗^), averaged across trials. In models 3 & 4, weights shown in crimson are estimated by linear regression to minimize the squared error between the activity of the OC units (𝐫𝐫_𝑡_) and the relative target location (𝐳̃_𝑡_ = 𝐳^∗^ − 𝐳_𝑡_), averaged across trials. Number of tuned parameters – 200, 10200, 400 and 600 for models 1, 2, 3 and 4 respectively. **(C) Left**: Correlation between gaze position (activity of the OC units) and the relative target position across test trials. **Middle**: Model performance in the test trials, quantified as variance in target position explained by stopping position (R-squared). **Right**: Correlation between target tracking error (TTE) and stopping error (SE). In all panels in C, *violin plots* represent distributions of statistics across 50 realizations of each model type and *cyan horizontal lines* denote average statistics from data.

By construction, only the models constrained by the behavioral strategy explain participants’ eye movements (Fig. 4C **– left**). Of these two models, only the one with tuned feedback from OC to FPN had good task performance (Fig. 4C **– middle; Suppl.** Fig. 3A, B). Notably, the performance of this model (**Model 4**) was almost as good as the task-optimized model in which all recurrent weights are tuned (**Model 2**) despite having substantially fewer tunable parameters (see figure caption). This suggests that recurrent frontoparietal-oculomotor (FPN-OC) interactions serve as a useful anatomical motif to support task performance. By selectively funneling the subspace of FPN activity that encodes the latent state (relative target position) into OC, Model 4 enables learning efficiently (*i.e*., with fewer tunable parameters) thereby highlighting the computational significance of embodied cognition. Unlike other models, Model 4 also recapitulates the trial-by-trial correlations that arise between target tracking error (TTE) and stopping error (SE) (Fig. 4C **– right**). This is because, in this model, error in estimating position results in poor target-tracking by the OC units that control eye movements, and this error propagates to joystick movements via tuned feedback connections from OC to FPN.

To test whether model 4 recapitulates more granular aspects of the data, we tested this model under two different levels of observation noise that differed by an order of magnitude (to simulate visual and inertial conditions) and found that the performance generalized to settings with greater observation noise albeit at a lower precision (Fig. 5A; compare with Fig. 1D). At the same time, the activity of the OC units in the model recapitulated the dynamics of eye movements seen in experiments (Fig. 5B; compare with Fig. 2A; **Suppl.** Fig. 3C). Specifically, the two OC units appeared to track the x-y components of the relative target position, regardless of the magnitude of observation noise. However, it is impossible to precisely discern the actual target position at any given time due to accumulation of noise. Consequently, the influence of target position on the OC network activity decreased during navigation. On the other hand, the influence of the stopping position increased, consistent with our experimental findings (Fig. 5C; compare with Fig. 4B). This suggests that the OC network encodes an internal estimate of the target location (i.e., belief), which is then used by FPN to control steering. Consistent with this interpretation, navigation performance deteriorated substantially when we prevented ‘eye movements’ in the model by clamping the activity of OC units to zero (Fig. 5D**, left**), which agrees with human performance when eye movements are inhibited (Lakshminarasimhan et al., 2020; Fig. 5D**, right**). Finally, we tested whether distributing the computation across the frontoparietal network in this manner enables the network to learn representations that resemble the brain. We previously showed that it is possible to dynamically decode target distance from population activity in monkey PPC (Lakshminarasimhan et al., 2023, Fig. 5E – **top**). We first verified that target distance can also be decoded from the model FPN activity (Fig. 5E – **middle**). Next, we reanalyzed the data to determine the subspace of PPC that is most informative about target distance. To do this, we first denoised the data by reducing it to the top 16 principal components. In this denoised subspace, we found that more than 90% of explainable variance was contained within the top five principal components (PC) of PPC activity (Fig. 5E – **bottom**, blue). Strikingly, target distance information was largely contained within the top few PCs of the strategy-constrained model (Fig. 5E – **bottom**, black solid) but not in the purely task-optimized model (Fig. 5E – **bottom**, black dashed). This is because, by explicitly projecting the belief about the relative target position into a low-dimensional OC activity, model 4 allows this signal to undergo recurrent amplification which increases its variance. In contrast, such amplification does not take place in the model lacking cognitive constraints (Model 2) where belief signals remain buried in low-variance modes (i.e., lower PCs) of the population activity.

**Figure 5:**
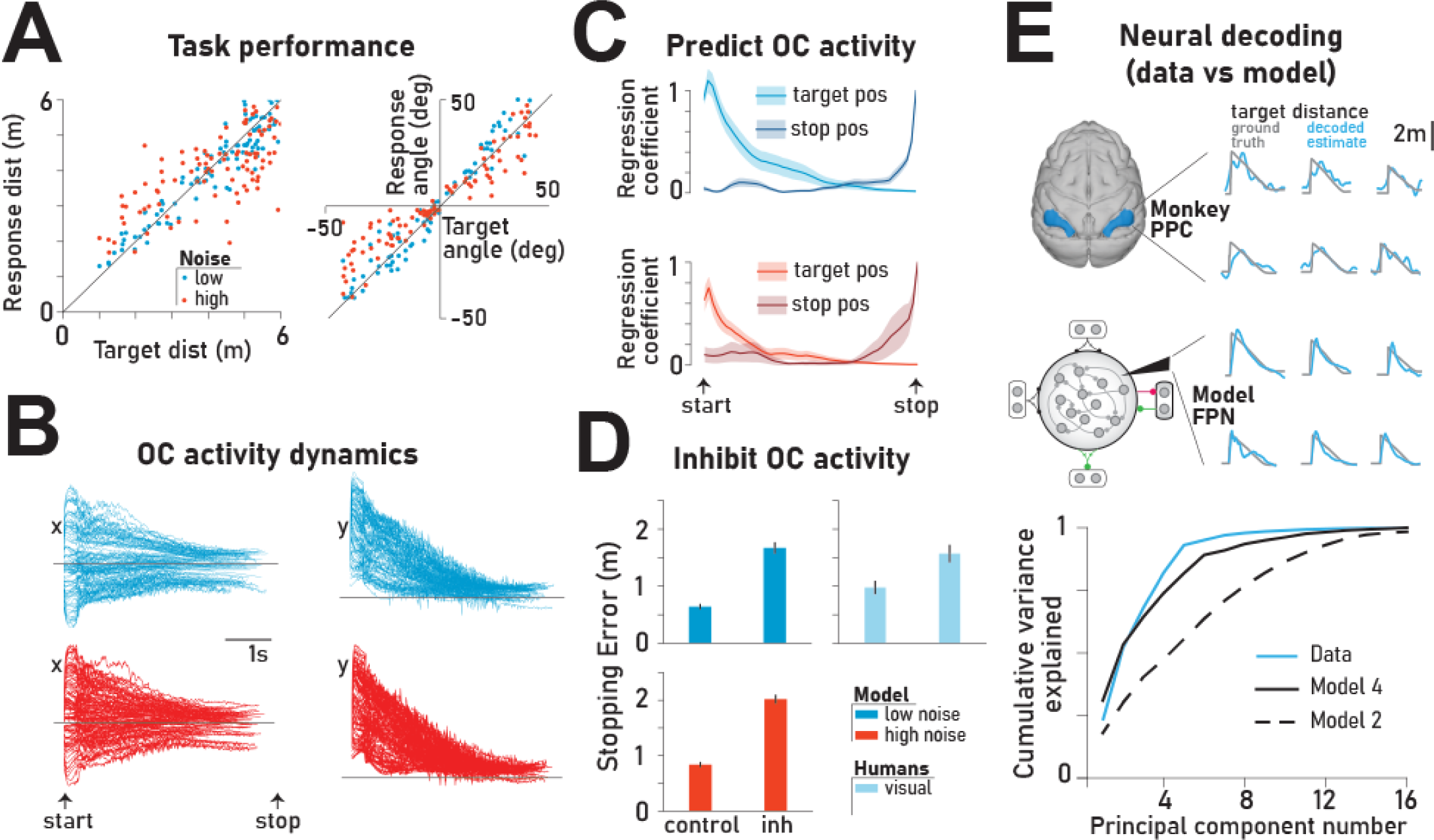
Strategy-constrained circuit model (Model 4) recapitulates behavioral and neural data. **(A)** Comparison of the radial distance (from origin) of the model’s stopping position against the radial distance of the target (**left**), as well as the angular eccentricity of the model’s stopping position versus target angle (**right**) across all trials. *Black lines* have unit slope. Visual (cyan) and vestibular (red) conditions are simulated by changing the variance of observation noise during testing (see Methods). **(B)** Activity dynamics of the two OC units that were constrained to encode the x and y components of the relative target position in a subset of test trials. Horizontal line denotes zero. **(C)** Coefficients corresponding to the target and stopping position from the linear regression model that best explained the OC unit activity (averaged across the two units). Error bars denote standard errors estimated by bootstrapping. **(D)** Error in stopping position, on typical trials (control) and trials in which the model OC units are inhibited by clamping their activity at zero (*inh*). For comparison, stopping error in free-gazing (control) and inhibited gaze (inh) trials in humans are shown on the right. Error bars denote standard errors estimated by bootstrapping. **(E)**. **Top**: Example trials showing the performance of a linear decoder trained to estimate target distance from a population of simultaneously recorded neurons in monkey PPC (average from 3 monkeys). **Middle**: Like the top panel but decoded from the model FPN activity. **Bottom**: Variance explained by the decoder as a function of the number of leading principal components used for decoding. Traces for models 1 & 3 (not shown) resemble models 2 & 4, respectively.

### Model Predictions

To study the implications of the embodied strategy, we simulated the effect of stimulating oculomotor units during the trial by systematically injecting a brief (0.2s) external input pulse of fixed amplitude that was either *positive* or *negative* into the model OC unit that encoded either the *horizontal* or *vertical* component of the believed target position, yielding four types of perturbations (right / left / up / down). The stimulation was delivered at various intervals following the removal of target position information (0s, 0.4s, 0.8s, 1.2s, 1.6s). Ideally, these perturbations should produce stereotyped rightward, leftward, upward, or downward saccades. In contrast, we noticed substantial variability in the evoked saccades regardless of the timing of the stimulation (Fig. 6A; **Suppl.** Fig. 4A). This variability could not be simply attributed to OC units participating in recurrent dynamics with FPN: Variability in saccades evoked was substantially lower when simulating the same stimulation protocol on the architecturally identical model lacking cognitive constraints (Model 2, Fig. 6B; **Suppl.** Fig. 4B).

**Figure 6:**
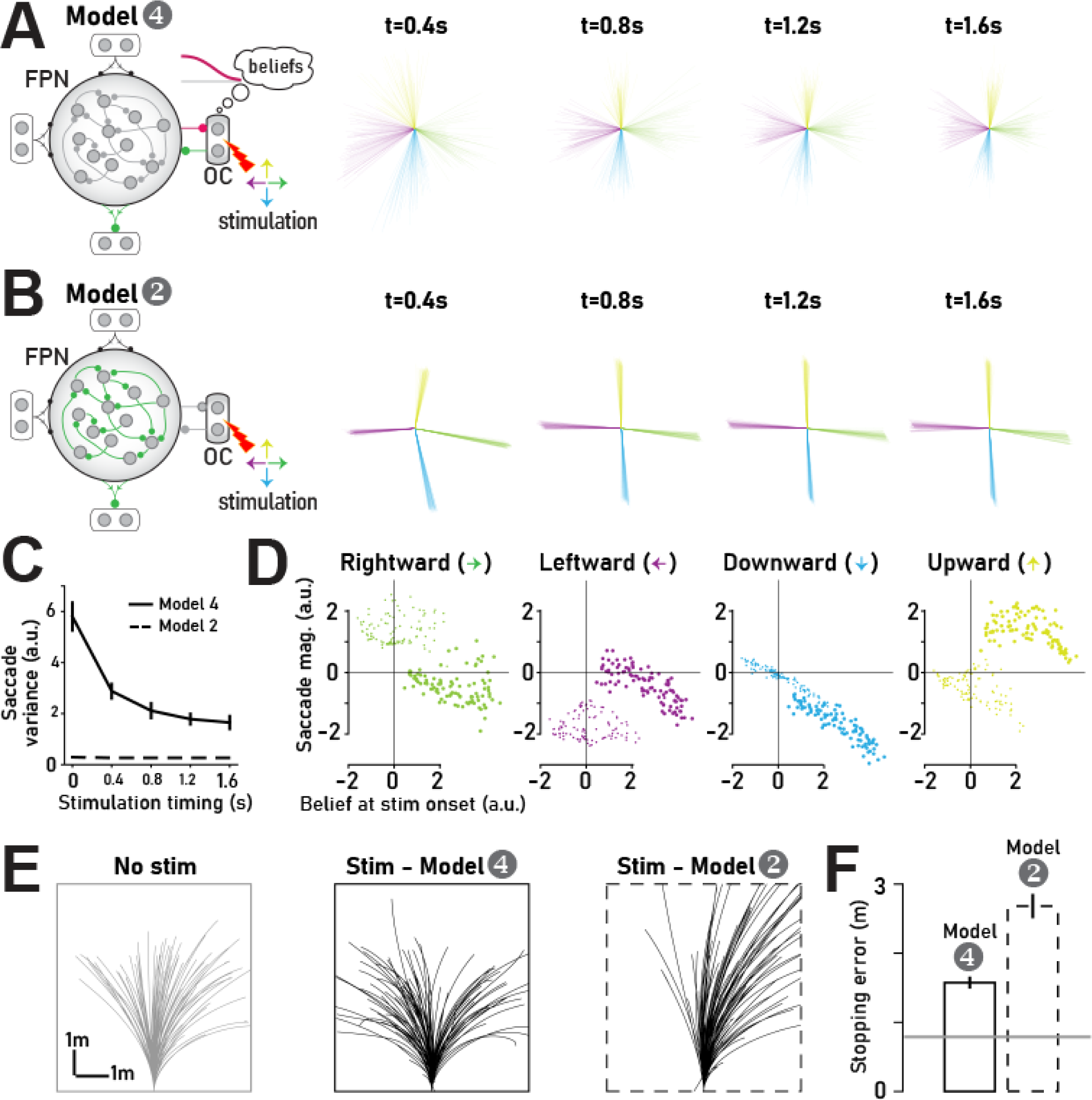
Model predictions for stimulation experiments. **(A)** Saccades evoked by stimulating the OC units at different delays with respect to the time at which the target position cue disappeared, while model 4 performed the navigation task. Each line corresponds to a different trial and color denotes the type of stimulation (see text). **(B)** Similar to A but showing the response of model 2. **(C)** Variability in the final position of the saccade across trials as a function of the timing of the stimulation. **(D)** Comparison of the magnitude of saccade against the model’s belief about the relative target position at the time of stimulation for 4 different stimulation sites (generating rightward, leftward, downward or upward saccades). *Thin* and *thick circles* denote the x (horizontal) and y (vertical) eye movement components respectively. Data plotted are from all stimulation times. **(E)** Movement trajectories of the model under baseline trials (left) and during stimulated trials (middle). Right: Trajectory of model 2 during stimulation. **(F)** Stopping errors of model 4 (solid) and model 2 (dashed) in stimulated trials. *Gray line* denotes the error during baseline trials. Error bars denote ±1 standard error of mean estimated by bootstrapping.

Furthermore, the variability of the evoked saccade gradually decreases as the stimulation occurred later in the trial (Fig. 6C). Since the variability in the belief (about the relative target position) also decreases as the trial progresses (see Fig. 5B), we asked whether the high variability in evoked saccade in the model was due to the trial-by-trial variability in beliefs. We found that the saccade magnitude was indeed strongly anti-correlated with the belief at the time of stimulation (mean Pearson’s *ρ* corr. coefficient ± SD across all conditions: x-component: −0.39 ± 0.2, y-component: −0.60 ± 0.1; Fig. 6D) suggesting that the belief substantially influences the properties of the evoked saccade. The negative sign in the correlation can be understood by recognizing that, for accurate navigation, the belief about the relative target position always approaches zero (recall Fig. 5B). Thus, saccades that are congruent with belief updates should be negative or positive depending on whether the belief is above or below zero, respectively. Furthermore, because the model encodes the x-y components of the beliefs in the two-dimensional eye position, memory about one component can persist even if the other component is perturbed by stimulation. For example, a stimulation intended to evoke a downward saccade might produce a saccade that is biased rightward or leftward depending on the horizontal component of the belief at the time of stimulation. Consequently, stimulation does not completely disrupt belief updates when the beliefs are embodied in eye movements. Indeed, the effect of stimulation on navigation performance (quantified by stopping errors) is relatively small in this model compared to the model that does not rely on the embodied strategy where stimulation can disrupt both components of the belief (model 2, Fig. 6E**-F**). Therefore, the model predicts that the embodied strategy should lead to a paradoxical effect wherein stimulating the oculomotor areas should evoke highly variable saccades, yet only modestly affect task performance.

## Discussion

Using a naturalistic behavioral paradigm defined by dynamic action/perception loops and unconstrained eye movements, we show that the dynamic belief about goal location is reflected in the subjects’ oculomotor behavior. By demonstrating that goal tracking is also observed in a purely inertial navigation version of the task in the absence of optic flow, we showed that this behavioral strategy is not driven by an active sensing strategy and instead provides strong support for the embodiment hypothesis. Specifically, we also show that these task-relevant eye movements reflect an embodiment of the *subjects’ dynamically evolving internal beliefs about the goal*, and not just the initial location of the target. Furthermore, we found that a neural model constrained by the cognitive strategy adopted by animals explains behavioral and neural data better than purely task-optimized models. Thus, we conclude that computations needed for steering are distributed across frontoparietal and oculomotor networks, where frontoparietal network temporally integrates self-motion but outsources belief state representation to the oculomotor network resulting in eye movements that dynamically track beliefs. We propose that mixing of signals between association and (oculo-)motor areas results from a distributed brain architecture which evolved to implement computations by grounding subjective beliefs about latent world states in states of the body.

### Internal belief embodiment hypothesis

We show for the first time that humans persistently use their eyes to track latent goal locations, even in the absence of visual navigational cues. In our previous study (Lakshminarasimhan 2020), we showed that the eyes follow the latent target in a visual-only condition where trial durations were much smaller (∼2 seconds), while inhibiting these eye movements worsened performance significantly, highlighting their computational importance. Here, we show that target-tracking can happen for much longer trial durations (>8 seconds), even in the absence of visual stimuli (inertial condition), and despite the presence of reflexive oculomotor processes (i.e., VOR). Specifically, we showed that target-tracking error was kept low for most of the trial in both visual and inertial conditions, although the error increased faster for the latter. We were able to map tracking errors to steering errors, where the correlation between these two quantities was higher later within a trial. Overall, these findings show that the eyes follow the believed goal location which shifts over time, from the actual location of the target when first presented, to the final stopping location. Eye movements have been found to facilitate working memory computations in non-navigation settings (Ballard et al., 1995; Johansson et al., 2012; Johansson & Johansson, 2014; Spivey & Geng, 2001) and other discrete domains (Gold & Shadlen, 2000; Loetscher et al., 2010; Tanenhaus et al., 1995). Our findings complementarily expand this body of work on embodied cognition to a naturalistic sequential decision task like navigation.

We generated a belief estimate as a dynamic weighted sum of the relative target and stopping positions, whose weights exhibit an almost perfect reversal between start and end of trial. This reconstructed belief modulates the saccades’ amplitude and direction, which proved crucial in the Inertial condition, as they allowed the eyes to successfully counter the VOR and track the believed goal location. VOR cancellation has been previously studied using targets that participants were required to fixate during passive yaw rotations (Crane & Demer, 1999; Lisberger, 1990). Here, we present the first evidence of volitional target-tracking eye movements countering the VOR in a naturalistic navigation setting. Importantly, these eye movements were driven dynamically by the belief about the relative goal location as participants actively steered toward it.

These experimental results provide strong support for the *embodiment* hypothesis, whereby by allowing dynamic beliefs about the relative target location to continuously modulate eye movements, the brain piggybacks on the oculomotor circuit and reduces the computational burden on working memory. This discovery was made possible by using a naturalistic behavioral paradigm, as opposed to highly-controlled tasks that restrict motor behavior and hinder the ability of the brain to use the algorithms that generate natural behaviors.

### Going beyond task-optimized computational models

Motivated by the support that our findings offer for the embodiment hypothesis as a strategy for navigational control, we propose a recurrent neural network (RNN) model of the underlying computation that the brain uses to exploit eye movements: a circuit model in which the believed target location is encoded in oculomotor neurons that have tuned bidirectional connections with the frontoparietal cortex (FPN) that integrates self-motion signals. This model with substantially fewer tuned connections was able to perform similarly to a model in which learning was accomplished by tuning all recurrent connections within the FPN. Notably, in addition to performing the steering task accurately, this model recapitulated human eye movements thereby providing a normative explanation for why subjective beliefs are externalized in eye movements. In contrast to purely task-optimized models, this strategy-constrained model also correctly predicted that leading principal components of the monkey posterior parietal cortex activity should encode their position relative to the goal.

The ability to predict neural responses accurately has made task-optimized neural network models an increasingly common tool for probing neural mechanisms underlying a wide range of computations including image recognition, speech perception, working memory, and motor control (Goehring et al., 2019; Martin Schrimpf et al., 2020; Martinez et al., 2017; Mi et al., 2017). However, such an approach neither explains why computations are distributed across functionally distinct modules nor allows modularity to emerge on its own. Our findings directly address this dual challenge by providing a possible computational benefit: both can be explained by augmenting task-optimized models with constraints obtained by analyzing the strategy used by animals to solve the task. Since naturalistic tasks increase the likelihood of engaging strategies that the brain evolved to use in the real world, we believe combining such task designs with strategy-constrained computational modeling can shed further light into distributed neural computations in other domains.

Note that we have focused here on FPN because neurons in the monkey posterior parietal cortex and dorsolateral prefrontal cortex have been identified as candidate regions involved in computing beliefs during this task (Noel et al., 2022; Lakshminarasimhan et al., 2023). Anatomical studies in monkeys have found extensive reciprocal connectivity between frontoparietal brain regions and neural circuits involved in eye movements, including frontal eye fields, supplemental eye fields, and area 8ar (Barbas & Mesulam, 1981; Huerta & Kaas, 1990; Leichnetz, 2001; Stanton et al., 1995).

For ease of interpretability, we have considered a minimal model of the oculomotor (OC) module with only two units. However, similar results could also be obtained by modeling the OC module as another RNN with a 2-dimensional output that controls horizontal and vertical eye position following previous work (H. Seung et al., 2000; H. S. Seung, 1996). Such an expanded model would still account for the amplification of belief signals seen in the monkey neural data as long as the neural activity in the OC module is low-dimensional. Furthermore, the computational benefit of learning FPN-OC interactions (over recurrent weights within FPN) will also hold provided the OC module has fewer units than FPN.

The model makes two concrete predictions to be tested in future experiments. First, the communication subspace between the FPN and oculomotor regions should represent the subjective beliefs about the relative position of the target. Second, stimulation of the oculomotor regions that provide feedback to FPN should have a modest yet clear effect on navigation performance. Regions with bidirectional connectivity with the posterior parietal and dorsolateral prefrontal cortex such as area 8ar, Frontal Eye Fields (FEF), Supplementary Eye Fields (SEF) (Barbas & Mesulam, 1981; Chavis & Pandya, 1976; Huerta & Kaas, 1990; Leichnetz, 2001; Stanton et al., 1995) are all excellent candidates for testing these predictions. More broadly, the proposed circuit model suggests that embodied cognition might be the reflection of a strategy by which the brain exploits distributed neural circuits and sensorimotor pathways structured through evolution in order to learn efficiently.

### Relation to previous works

Our emphasis of the role of eye movements in dynamically tracking latent beliefs complement previous studies that highlight the information-gathering role of temporally structured eye movements (Ahmad et al., 2014; Hoppe & Rothkopf, 2016, 2019; S. C.-H. Yang et al., 2016; Zhu et al., 2022) and contextualize findings from controlled studies that report an influence of short-term memory on smooth pursuit eye movements (Adams et al., 2015a; Deravet et al., 2018; Jean-Jacques Orban de Xivry et al., 2013).

The proposed model builds on recent efforts that take advantage of well-characterized behavioral strategies to gain mechanistic insights via neural network models. For example, Molano-Mazón et al., (2023) demonstrated a need to incorporate structural priors into RNNs (via pre-training) for recapitulating suboptimal choice by rats that fail to account for serial correlations in stimulus statistics across trials. Likewise, Andalman et al., (2019) varied interaction strengths in an RNN model to account for a stress-induced switch from active to passive coping strategy in zebrafish. In another recent study, Rajalingham et al., (2022) endowed RNNs with an auxiliary loss function to mimic human error patterns in an intuitive physics task. However, to our knowledge, no study has harnessed a dynamic, within-trial behavioral strategy to inform the design of such models nor shown the need to use modular architectures to replicate animal behavior. The present study achieves both by using a naturalistic task to tap into an innate, evolutionarily conserved behavioral strategy for tracking one’s beliefs over time.

Recent work has contributed statistical tools to infer latent beliefs from behavior (Adams et al., 2015b; Kumar et al., 2019; Lee et al., 2014; Reddy et al., 2018; Sohn et al., 2019); our findings and proposed model could facilitate the development and application of these tools in sequential decision behaviors. Although our model is simplistic, it can guide future studies that probe neural mechanisms underlying the involvement of the oculomotor system in cognition. Also, the learning efficiency of the distributed architecture has important implications for realizing biologically inspired artificial intelligence in embodied agents, especially robotics.

Embodiment and its computational role in cognition have been largely overlooked by the neuroscience community, and yet their importance on artificial agents is the subject of an ongoing debate with decades-long roots (Abdou et al., 2021; Bender & Koller, 2020; Chalmers, 2023; Harnad, 1990; Lake & Murphy, 2023; LeCun, 2022; Piantadosi & Hill, 2022). Our study underlines embodiment as a cornerstone of human intelligence that any attempt to human-like computations and representations in machines should seriously consider.

## Methods

### EXPERIMENTAL MODEL AND SUBJECT DETAILS

8 subjects (6 Male, 2 Female; all adults in the age group 18-32) participated in the eye-tracking experiments. Apart from two subjects, all subjects were unaware of the purpose of the study. Experiments were first performed in the above two subjects before testing others. All experimental procedures were approved by the Institutional Review Board at Baylor College of Medicine and all subjects signed an approved consent form.

### METHOD DETAILS

#### Behavioral task – Visual, Inertial and Multisensory motion cues

The task required subjects to navigate to a remembered location on a horizontal virtual plane using a joystick, rendered in 3D from a forward-facing vantage point above the plane. Visual and/or vestibular sensory feedback was provided. Visual feedback was stereoscopic, composed of flashing triangles to provide self-motion information but no landmark. Vestibular feedback was generated by a moving platform approximating the properties of their virtual self-motion.

Participants pressed a button on the joystick to initiate each trial and were tasked with steering to a randomly placed target that was cued briefly at the beginning of the trial. A short tone at every button push indicated the beginning of the trial and the appearance of the target. After one second, the target disappeared, which was a cue for the subject to start steering. Participants were instructed to stop at the remembered target location, and then push the button to register their final position and start the next trial. Participants did not receive any feedback about their performance. Prior to the first session, all participants performed about ten practice trials to familiarize themselves with joystick movements and the task structure.

Participants performed the task under three sensory conditions, which were interleaved randomly across trials. In the visual condition, participants had to navigate towards the remembered target position given only visual information (optic flow); no vestibular sensory feedback was provided during motion. In the combined condition, subjects were provided with both visual and inertial (vestibular/somatosensory) information during their movement. In the Inertial condition, after the target disappeared, the entire visual stimulus was shut off too, leaving the subjects to navigate in complete darkness using only inertial cues.

Independently of the manipulation of the sensory information, the properties of the motion controller also varied from trial to trial. Participants experienced different time constants in each trial, which affected the type and amount of control that was required to complete the task. In trials with short time constants, joystick position mainly controlled velocity, whereas in trials with long time constants, joystick position approximately controlled the acceleration (explained in detail in *Control Dynamics* Methods section in Stavropoulos et al., 2022).

Each participant performed a total of about 1450 trials (mean ± standard deviation (SD): 1450 ± 224), split equally among the three sensory conditions (mean ± SD – vestibular: 476 ± 71, visual: 487 ± 77, combined: 487 ± 77).

#### Visual stimulus

The virtual world comprised a ground plane whose textural elements whose lifetimes were limited (∼250ms) to avoid serving as landmarks. The ground plane was circular with a radius of 37.5m (near and far clipping planes at 5cm and 3750cm respectively), with the subject positioned at its center at the beginning of each trial. Each texture element was an isosceles triangle (base × height 5.95 × 12.95 cm) that was randomly repositioned and reoriented at the end of its lifetime. The floor density was held constant across trials at 𝜌 = 2.5 elements⁄m^2^. The target, a circle of radius 25cm whose luminance was matched to the texture elements, flickered at 5Hz and appeared at a random location between 𝜃 = ±38° of visual angle at a distance of 𝑟 = 2.5 − 5.5 m (average distance 𝑟̅ = 4 m) relative to where the participant was stationed at the beginning of the trial. The stereoscopic visual stimulus was rendered in an alternate frame sequencing format and subjects wore active-shutter 3D goggles to view the stimulus.

#### Experimental setup

The participants sat comfortably on a chair mounted on an electric motor allowing unrestricted yaw rotation (Kollmorgen motor DH142M-13-1320), itself mounted on a six-degree-of-freedom motion platform (comprised of MOOG 6DOF2000E). Subjects used an analog joystick (M20U9T-N82, CTI electronics) with two degrees of freedom and a circular displacement boundary to control their linear and angular speed in a virtual environment based on visual and inertial stimuli. The visual stimulus was projected (Canon LV-8235 UST Multimedia Projector) onto a large rectangular screen (width × height: 158 × 94 cm) positioned in front of the subject (77 cm from the rear of the head). Participants wore crosstalk-free ferroelectric active-shutter 3D goggles (RealD CE4s) to view the stimulus. Participants wore headphones generating white noise to mask the auditory motion cues. The participants’ head was fixed on the chair using an adjustable CIVCO FirmFit Thermoplastic face mask. Eye movements were monitored at 120Hz using ISCAN 06-604-0302 binocular eye-tracking and ISCAN ETL 500 software.

#### Joystick control

Participants navigated in the virtual environment using a joystick placed in front of the participant’s midline, in a holder mounted on the bottom of the screen. This ensured that the joystick was parallel to the participant’s vertical axis, and its horizontal orientation aligned to the forward movement axis. The joystick had two degrees of freedom that controlled linear and angular motion. Joystick displacements were physically bounded to lie within a disk, and digitally bounded to lie within a square. Displacement of the joystick over the anterior-posterior (AP) axis resulted in forward or backward translational motion, whereas displacement in the left-right (LR) axis resulted in rotational motion. The joystick was enabled after the disappearance of the target. To avoid skipping trials and abrupt stops, the button used to initiate trials was activated only when the participant’s velocity dropped below 1 cm/s.

The joystick controlled both the visual and inertial stimuli through an algorithm that involved two processes. The first varied the <u>control dynamics (CD)</u>, producing velocities given by a leaky integration of the joystick input, mimicking an inertial body under viscous damping. The time constant of the leak (leak constant) was varied from trial to trial, according to a random walk (**Suppl. Fig S2**).

The second process was a <u>motion cueing (MC)</u> algorithm applied to the output of the CD process, which defined physical motion that approximated the accelerations an observer would feel under the desired control dynamics, while avoiding the hardwired constraints of the motion platform. This motion cueing algorithm trades translation for tilt, allowing extended acceleration without hitting the displacement limits of the platform.

These two processes are explained in detail in Stavropoulos et al., 2022.

#### Stimulus and data acquisition

All stimuli were generated and rendered using C++ Open Graphics Library (OpenGL) by continuously repositioning the camera based on joystick inputs to update the visual scene at 60 Hz. The camera was positioned at a height of 70cm above the ground plane. Spike2 software (Power 1401 MkII data acquisition system from Cambridge Electronic Design Ltd.) was used to record and store the target location (𝑟, 𝜃), subject’s position (𝑟̃, 𝜃̃), horizontal positions of left and right eyes (𝛼_𝑙_ and 𝛼_𝑟_), vertical eye positions (𝛽 and 𝛽) and all event markers for offline analysis at a sampling rate of 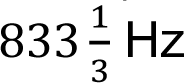.

## QUANTIFICATION AND STATISTICAL ANALYSIS

Customized MATLAB code was written to analyze data and to fit models. Depending on the quantity estimated, we report statistical dispersions either using 95% confidence interval, standard deviation, or standard error in the mean. The specific dispersion measure is identified in the portion of the text accompanying the estimates. For error bars in figures, we provide this information in the caption of the corresponding figure. We report and describe the outcome as significant if 𝑝 < 0.05.

### Bias estimation

In each sensory condition, we first computed the τ-independent bias for each subject; we regressed (without an intercept term) each subject’s response positions (𝑟̃, 𝜃̃) against target positions (𝑟, 𝜃) separately for the radial (𝑟̃ vs 𝑟) and angular (𝜃̃vs 𝜃) coordinates, and the radial and angular multiplicative biases were quantified as the slope of the respective regressions (**Fig 2A**). In addition, we followed the same process to calculate bias terms within three τ groups of equal size (Fig. 2C).

### Characterizing eye, target and stop position in eye coordinates

For convenience, we express the subject’s *actual* eye position using the following two standard degrees of freedom: (𝑖𝑖) Conjunctive horizontal movement of the two eyes quantified here as the mean lateral position of the two eyes, 𝛼 = (𝛼_left_ + 𝛼_right_)⁄2, (𝑖𝑖𝑖𝑖) Conjunctive vertical movement of the two eyes quantified here as 𝛽 = (𝛽_left_ + 𝛽_right_)⁄2. Disjunctive horizontal and vertical eye movements (horizontal and vertical vergence, respectively) were not considered for our analysis, because of the documented difficulty in humans to execute vergence to imagined moving objects (Erkelens et al., 1989; Lakshminarasimhan et al., 2020).

To test whether participants’ eyes tracked the location of the (invisible) target, we need the target and eye positions to be on the same reference frame. Therefore, we transformed target position from world to eye coordinates. Let *s* denote the stage of trial evolution, i.e. percentage of total distance travelled from 0 to 100%. We denote the target position in world coordinates as (𝑥_t_, 𝑦_t_, 𝑧_t_), relative to the midpoint of the participant’s eyes at trial stage *s*. The target position in eye coordinates - relative to fixating at the point (0,0, ∞) - relates to its position in world coordinates as:

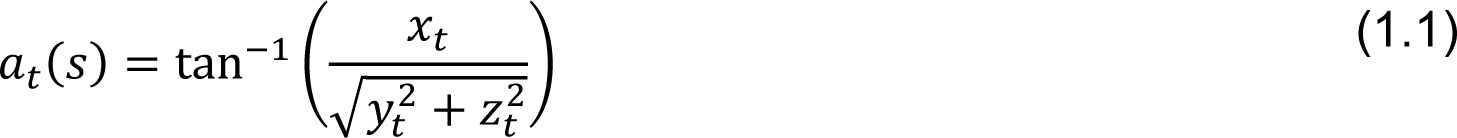

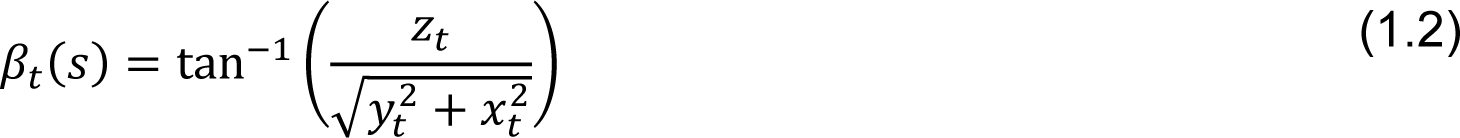

Where 𝑎_𝑡_(𝑠) and 𝛽_𝑡_(𝑠) are horizontal and vertical target positions at trial stage *s*, respectively. Note that 𝑧_𝑡_ is determined only by the viewing height, and therefore remains constant. On the contrary, 𝑥_𝑡_, 𝑦_𝑡_ change continuously as the participants steer in the virtual environment.

In approximately 8% of the trials, the subject travelled beyond the target. The target position in eye coordinates towards the end of these trials was outside the physical range of gaze. Therefore, we removed time points at which any of the two components of target position in **Equation 1** exceeded 60° before further analysis. Such time points constituted less than 1% of the dataset and including them did not qualitatively alter the results.

Similarly, we calculated 𝑎_𝑠_(𝑠) and 𝛽_𝑠_(𝑠) as the horizontal and vertical stopping positions in eye coordinates.

### Target-tracking error and belief analysis

We tested how target-tracking performance was associated with steering performance, by estimating the correlation between steering and target-tracking errors across trials (Fig. 2C, D). As mentioned above, we scaled trials according to the percentage of total distance travelled and computed this correlation as trials evolved. At every trial stage, *steering error* was given by the Euclidean distance between the target and stop positions in eye coordinates as 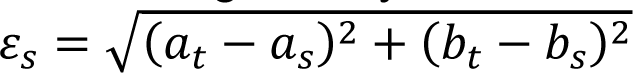, while *target-tracking error* (TTE) was given by the Euclidean distance between eye and target position as 𝜀_𝑡_ = 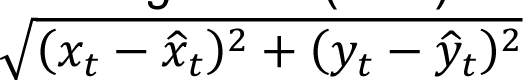, where 𝜀_𝑡_ is the TTE, and (𝑥_𝑡_, 𝑦_𝑡_) and (𝑥)_𝑡_, 𝑦)_𝑡_) are the horizontal and vertical coordinates of the target and eye position, respectively. Chance level TTE was estimated as the mean of the null distribution obtained by shuffling target positions across trials. The same method was used to calculate the correlation between steering and tracking errors for the *saccade-free eye movements* (discussed below).

To compute an estimate of the participants’ belief about the target location we regressed the participants’ eye position against the target and stop positions (multiple regression), obtaining a kernel of weights for each position over trial progression (Fig. 2E). As all trials are scaled equally this way, we regressed the eye positions at trial stage *s* against the corresponding target and stop positions for the horizontal and vertical components, separately. This provided us with regression weight kernels for the target and stop positions of each component from 0 to 100% of total distance travelled.

To reconstruct this belief, we simply multiplied the target and stop positions with their respective weights at each trial stage *s*, for each participant (Fig. 3C).

### Saccade analysis

For saccade detection, we estimated the instantaneous speed of eye movements as (𝛼̇ ^2^ + 𝛽^𝛽̇2^)^1/2^ where 𝛼 and 𝛽 denote horizontal and vertical eye positions respectively (as defined above), and the dot denotes a time derivative. Saccades were detected by identifying the time points at which the speed of eye movements crossed a threshold of 150°/s (a threshold of 25°/s yielded similar results). Specifically, saccade onset was detected as the time point at which the speed of eye movements crossed the threshold from below, and saccade offset as the time at which the speed dropped below the threshold. The amplitude of saccades was taken to be the average displacement of the position of the two eyes from saccade onset to 150ms later (Δ𝛗 = (Δ𝛼^2^ + Δ𝛽^2^)^1/2^).

To explore the contribution of saccades in target-tracking, we generated saccade-free eye movements by subtracting the displacement of the eye position caused by saccades after target offset (𝑡 ≥ 1𝑠) (Fig. 3A, B). We removed the periods between saccade onset and offset from the eye velocity signal. The remaining signal was linearly interpolated and then integrated to calculate eye displacement independent of saccades (saccade-free eye displacement). Finally, the eye position at the time of target offset was added to the saccade-free eye displacement. We then computed the correlation between steering and tracking errors for the *saccade-free eye movements* (just as we did for the actual eye position; see *Comparing steering and target-tracking errors*).

To test that the eyes would reflect target beliefs even under the control of the VOR, we explored the relationship between the cumulative saccade amplitude in each trial and the corresponding steering error (Fig. 3B). We only considered the horizontal component of saccades (Δ𝛼) which is aligned to the evoked VOR during rotation. Therefore, we estimated the Pearson’s correlation coefficient between angular steering errors and horizontal cumulative saccade amplitudes.

To quantify the precise relationship between saccade amplitude and tracking error (TTE), we obtained a regression weight kernel by regressing horizontal and vertical amplitudes of the saccade (Δ𝛼 and Δ𝛽) on horizontal and vertical target-tracking errors (𝛼 − 𝑎_𝑡_ and 𝛽 − 𝛽_𝑡_), respectively, at various lags between ±1𝑠 with 𝑙^2^regularization (Fig. 3C). Similarly, we computed the kernels for the stop position-tracking error (SPTE) and the belief-tracking error (BTE; based on reconstructed belief, see Methods: *Target-tracking error and belief analysis*).

Finally, we computed the gain of the eye position with respect to the target, to evaluate the effect of saccadic eye movements on target-tracking (Suppl. Fig. 2C). Specifically, we regressed (without intercept) the eye positions at time *t* against the corresponding target positions for the vertical and horizontal components, separately. We performed this regression for both the actual and the saccade-free eye positions.

### Recurrent neural network models

We trained four different recurrent neural network (RNN) models to solve the velocity control version of the task performed by human participants. All models comprise two modules: one recurrently connected population of 100 nonlinear (‘sigmoidal’) units that we identify as the frontoparietal circuit (FPN) and an oculomotor (OC) module comprising two linear units encoding vertical and horizontal eye position where for simplicity, we ignore the biomechanics of eye movement generation. The FPN module contains 4 input channels, two for conveying the 2D target location (𝐳^∗^) encoded in the amplitude of a transient pulse delivered in the beginning of the trial and two for conveying continuous sensory feedback about the 2D self-motion velocity (𝐳̇) throughout the trial. There were 2 output channels from FPN, one each for controlling the velocity of the ‘hand’ along the linear and angular axes of the joystick i.e., movement acceleration (ż). To mimic process noise, we added zero-mean additive gaussian noise to the output channels, and the noisy output is temporally integrated and fed back to the network through the input channels conveying movement velocity, thereby closing the sensorimotor loop. Additive noise is also added to the input channels to simulate observation noise. This feedback mimics the functionality of the virtual reality simulator that uses the joystick output to render real-time sensory feedback in the form of optic flow cues or vestibular cues in our experiments.

The equation governing the network dynamics was:

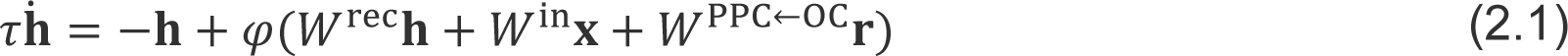

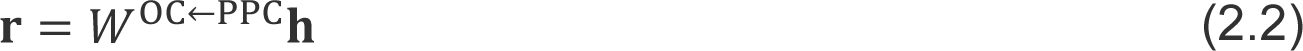

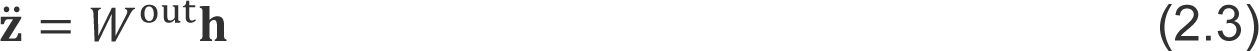

where **h** and **r** represent population activity in FPN and OC respectively. 𝐱 = (𝐳^)̇^, 𝐳^∗^) denotes the input to PPC where 𝐳^)̇^ = 𝐳̇ + 𝜺 denotes the velocity corrupted by additive observation noise 𝜀∼𝒩(0, 𝜎^2^). ż is the network output representing acceleration such that 𝐳̇ = ∫ d𝐳 𝐳̃ where 𝐳̃ = ż + 𝜂 denotes acceleration corrupted by process noise 𝜂∼𝒩(0, 𝜎^2^). 𝜏 is the cell-intrinsic time constant, and 𝜑(·) = tanh(·) is the neuronal nonlinearity. Matrices 𝑊^rec^, 𝑊^in^, 𝑊^out^, 𝑊^PPC←OC^ and 𝑊^OC←PPC^ correspond to recurrent, input, output, fronto-parietal and parieto-frontal weights respectively.

### Model training and details

We trained the RNN models defined in Eq. (2) by tuning different sets of model parameters in each model. Models were trained to reach the target location within a certain time *t*\* and stay there for 0.6s. *t*\* corresponded to the time taken when traveling along an idealized circular trajectory from the starting location to target location at maximum speed. The time-constant 𝜏 was set to 20 ms and each training trial lasted between 2-3 s depending on the target location. In all four models, output weights 𝑊^out^ were updated at the end of each trial to minimize the loss function 𝐿 = ∑_𝑡>𝑡_∗ |𝐳(𝑡) − 𝐳^∗^|^2^, using gradient descent. In addition to output weights, we updated the recurrent weights 𝑊^rec^ in Model 2 and feedback weights from OC to FPN (𝑊^FPN←OC^) in Model 4 to minimize 𝐿 via backpropagation through time. In models 3 and 4, behavioral strategy constraint was incorporated by training the weights from FPN to OC (𝑊^OC←FPN^) were trained to minimize an auxiliary loss function 𝐿_aux_ = ∑_𝑡_ |𝐫(𝑡) − 𝐳̃(𝑡)|^2^ by linear regression where the relative target location 𝐳̃(𝑡) = 𝐳(𝑡) − 𝐳^∗^(𝑡). Since abruptly updating 𝑊^OC←FPN^ in conjunction with other weights hampered learning, we updated 𝑊^OC←PPC^ incrementally as 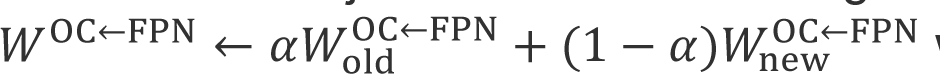 where 𝛼 = 0.99. The total number of free parameters were 200, 10 200, 400 and 600 for models 1, 2, 3 and 4 respectively.

### Neural recordings

Three rhesus macaques (*Macaca mulatta*) (all male, 7-8 years. old)—referred to as B, S, and Q for simplicity—participated in the experiments. All surgeries and experimental procedures were approved by the Institutional Review Board at Baylor College of Medicine, and were in accordance with National Institutes of Health guidelines.

Monkeys were chronically implanted with a lightweight polyacetal ring for head restraint, and scleral coils for monitoring eye movements (CNC Engineering, Seattle WA, USA). Utah arrays were chronically implanted in area 7a of the Posterior Parietal Cortex in the left hemisphere of all three monkeys using craniotomy. Prior to the surgery, the brain area was identified using structural MRI to guide the location of craniotomy. After craniotomy, the array was pneumatically inserted after confirming the coordinates of the target area using known anatomical landmarks.

At the beginning of each experimental session, monkeys were head-fixed and secured in a primate chair placed on top of a platform (Kollmorgen, Radford, VA, USA). All methods regarding these recordings have been previously described in Lakshminarasimhan et al., 2023.

## DATA AND SOFTWARE AVAILABILITY

MATLAB and Python code implementing all quantitative analyses and modelling, respectively, is available online (https://github.com/AkisStavropoulos/eye_movement_analysis and https://github.com/kaushik-l/firefly-rnn). Datasets generated by this study are available online (https://gin.g-node.org/akis_stavropoulos/belief_embodiment_through_eye_movements_facilitates_memory-guided_navigation).

## Acknowledgements

We thank Jing Lin and Jian Chen for their technical support. This work was supported by NIH grant 1R01 DC004260 and 1R01 NS127122 to D.E.A., NSF NeuroNex 1707400, NIH CRCNS 1R01 NS120407-01, 1U19 NS118246, and Simons Collaboration on the Global Brain, grant no. 324143 to D.E.A.

## Notes

https://gin.g-node.org/akis_stavropoulos/belief_embodiment_through_eye_movements_facilitates_memory-guided_navigation

### Competing Interest Statement

The authors have declared no competing interest.

