## Supplemental Figure 1 for "Belief embodiment through eye movements facilitates memory-guided navigation"

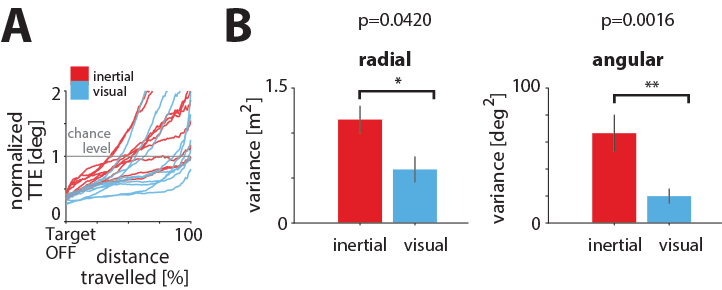

**Suppl. Figure 1: Additional performance measures. (A)** Normalized TTE over trial progression (percentage of total distance travelled), shown individually for each subject. TTE was normalized by the chance-level TTE obtained by shuffling (*gray line*). **(B)** Radial (left) and angular (right) variance of stopping locations, after subtracting the corresponding bias (see Fig. 1D; variance of residual errors). *Error bars* denote ±1 SEM. Notice the higher response variability in the inertial condition, presumably due to higher sensory uncertainty (Stavropoulos et al., 2022).
