## Supplemental Figure 2 for "Belief embodiment through eye movements facilitates memory-guided navigation"

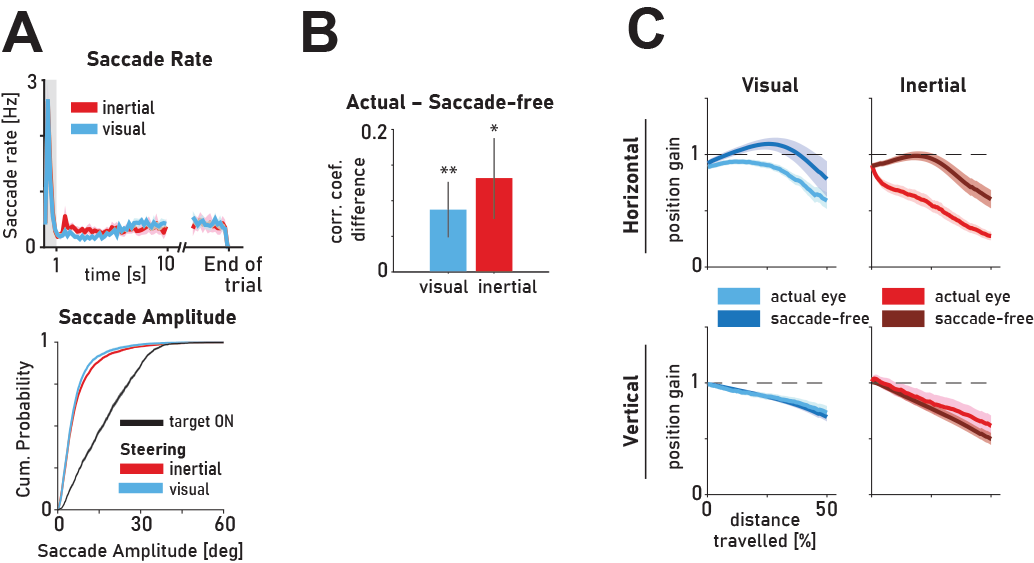

**Suppl. Fig. 2: (A)** **Top:** Time-course of the rate of saccades during the trial, averaged across all subjects. G*ray region:* target on period. **Bottom:** Cumulative distribution of saccade amplitude (within first 10s of each trial) conditioned on the task epoch (target presentation, steering), averaged across participants. **(B)** Difference between correlation coefficients (actual – saccade-free) from (A) for the horizontal components of TTE and SE. *Asterisks* denote the level of statistical significance of the correlation difference within each condition (*: p<0.05, **: p<0.01). Error bars: ±1 SEM. **(C)** Gain of eye position with respect to target position, with and without considering saccadic eye movements. Because horizontal slow eye movements are strongly affected by the yaw vestibulo-ocular reflex (VOR), which would be unaffected by internal beliefs, we reasoned that saccades would have a large contribution to the difference between the actual and believed target location. Indeed, in contrast to the actual eye position (which includes saccadic contribution), the gain between horizontal saccade-free eye position and actual target position (computed by regressing eye position against the corresponding target position at each percentile of distance travelled for either the actual or saccade-free eye positions) is close to 1, suggesting that the VOR drives the horizontal component of slow eye movements during yaw rotation.
