## Supplemental Figure 3 for "Belief embodiment through eye movements facilitates memory-guided navigation"

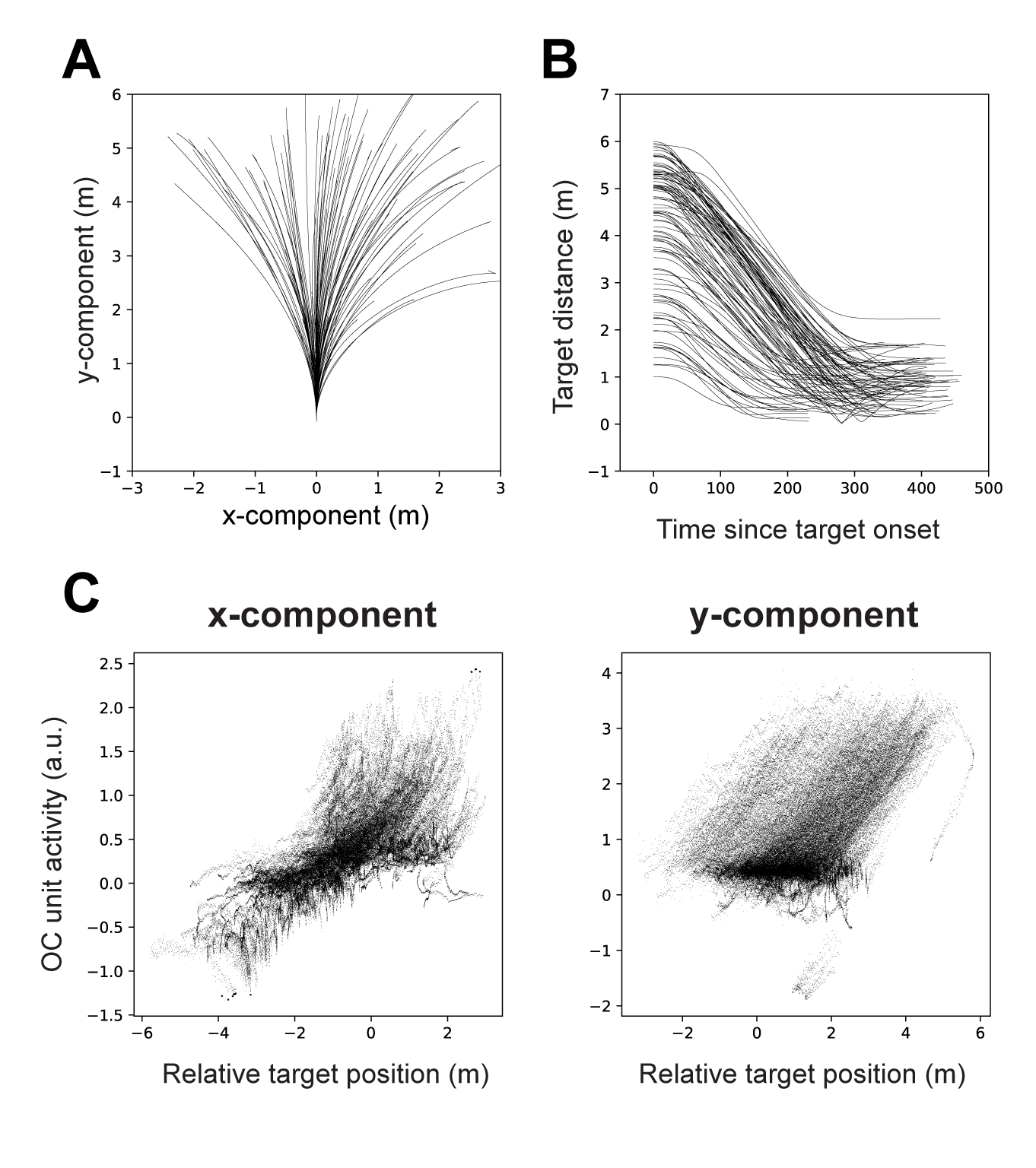

**Suppl.** **Fig. 3: Predictions of model 4. (A)** Example trajectories. **(B)** Evolution of target distance predicted by the model as a function of time. **(C)** Comparison of the activity of the units in the OC module against the relative target position across trials. Each dot corresponds to a single time point from one trial. Comparison is done separately for the x and y components, demonstrating a strong correlation in both components.
