## Supplemental Figure 4 for "Belief embodiment through eye movements facilitates memory-guided navigation"

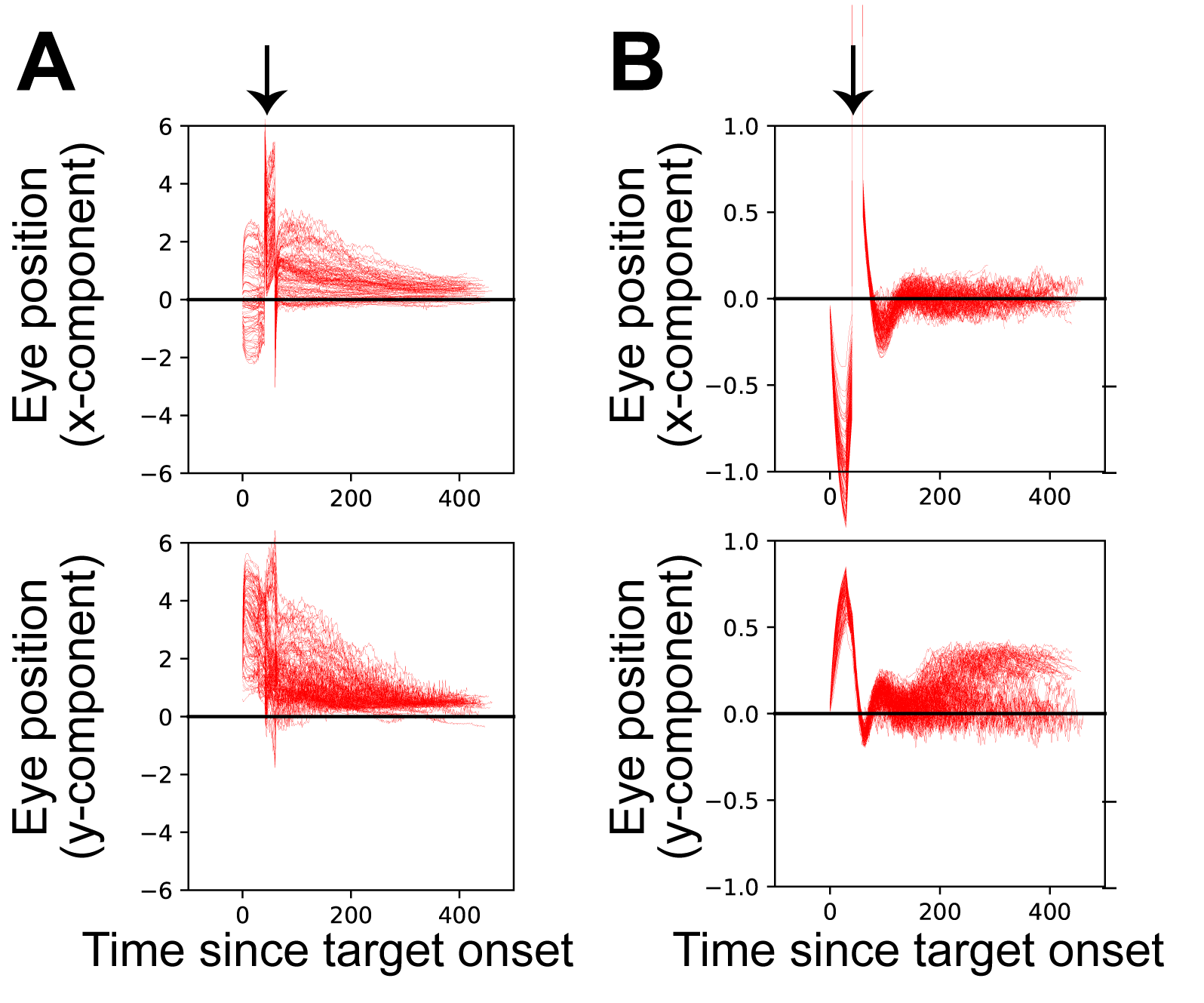

**Suppl.** **Fig. 4: Simulations of electrical stimulation on oculomotor (OC) units of models 4 and 2. (A)** **Top**: Activity dynamics of the x-component (**top**) and y-component (**bottom**) of eye position (Model 4). **(B)** Similar to A, but for the set of trials in which stimulation was delivered to model 2. *Arrow* indicates the time of stimulation delivery.
